# Estimation of contemporary effective size of large continuously distributed populations

**DOI:** 10.64898/2026.02.09.704747

**Authors:** Armando Caballero, Santiago C. González-Martínez, Enrique Santiago

**Affiliations:** Centro de Investigación Mariña, Universidade de Vigo, 36310 Vigo, Spain; INRAE, Univ. Bordeaux, BIOGECO, Cestas, France; Departamento de Biología Funcional, Facultad de Biología, Universidad de Oviedo, 33006 Oviedo, Spain

**Keywords:** Genetic drift, inbreeding, linkage disequilibrium, SNP, neighbourhood size, maritime pine

## Abstract

Estimation of the effective size (*N*_*e*_) of large populations with a continuous distribution across wide geographic areas and limited dispersal of individuals has been elusive so far. Estimates of the contemporary *N*_*e*_ from genetic markers for such large, structured populations, typically of plant and marine species, tend to be strongly biased downwards, which has led to question their relevance. Here we show that a recently proposed estimation method of *N*_*e*_ from linkage disequilibrium between markers, which accounts for population structure, yields estimates of metapopulation *N*_*e*_ when the sampling area is sufficiently large. The method is applied to empirical data of maritime pine (*Pinus pinaster* Aiton). While previous estimates of *N*_*e*_ in pine populations were of the order of a few hundred individuals, we show that estimates of the metapopulation *N*_*e*_ can reach values of the order of tens of thousands of individuals. This result is especially relevant from a conservation point of view, as populations with *N*_*e*_ lower than 500 individuals are considered to be under the risk of extinction.

## Introduction

The effective population size (*N*_*e*_) is a key metric in evolutionary biology, conservation of threatened species, and the breeding of domestic animals and crops, as it reflects the degree of genetic drift and inbreeding in a population (Wright 1931, Caballero 2020, Chap. 5). This statistic is widely considered as a key indicator for evaluating the conservation status of populations (Hoban et al. 2020, Clarke et al. 2024). There are various methods to estimate the *N*_*e*_ of a population from genetic markers, based on the change of allele frequencies or the linkage disequilibrium (LD) between loci, among other methods (Wang et al. 2016). Most methods of estimation of *N*_*e*_ assume that the population under study is a closed one without immigrants from other populations. However, a common situation is that of a metapopulation structured in different populations with migration between them. In this case, the estimates of *N*_*e*_ from genetic markers can produce biased estimates of both the local *N*_*e*_ and the metapopulation *N*_*e*_ (Waples and England 2011, Ryman et al. 2019, Novo et al. 2023, Kardos and Waples 2024). The software MLNe (Wang and Whitlock 2003, Wang 2022) provides estimates of *N*_*e*_ assuming population structure, but it is restricted to a model where a small focal population is considered a sink receiving migrants from an infinitely large source population. A recent method has been developed to estimate *N*_*e*_ which accounts for population structure under more general scenarios (Santiago et al. 2025). Based on an island model (Wright 1931), the method provides estimates of the metapopulation *N*_*e*_, the migration rate between subpopulations, the degree of genetic differentiation between them measured by Wright’s (1943) fixation index *F*_*ST*_, and the number of subpopulations.

A common scenario in plants and marine fishes, which however differs from the island model and other structure population models, occurs when large continuously distributed populations are extended over large areas featuring isolation by distance (IBD). In this situation, the neighbourhood size (*N*_*S*_) (Wright 1946) is a measure that describes the local area within which most matings occur. Under this scenario, if sampling is made in a local area close to the effective range of dispersal of the species, the estimates of *N*_*e*_ by LD or allele frequency changes over time of genetic markers, are close to the neighbourhood size (Neel et al. 2013, Nunney 2016). As the geographic scale of sampling is increased, the *N*_*e*_ estimates increase towards the global *N*_*e*_, but a plateau well below this value is reached, which underpins the effect of underestimation of *N*_*e*_ under population structure (Neel et al. 2013, Shirk and Cushman 2014, Waples 2024). This can be an explanation for the extremely low ratios of *N*_*e*_/*N* found for marine fish species (Marandel et al. 2019) and forest trees (Santos-del-Blanco et al. 2022), which questions the value of *N*_*e*_ estimated from genomic markers in conservation programmes (Fady and Bozzano 2021). For example, estimates of *N*_*e*_ for 19 continuously distributed populations of maritime pine (*Pinus pinaster* Aiton) obtained by Santos-del-Blanco et al. (2022) using LD between 3,514 SNPs provided values with an average *N*_*e*_ of 375 (range 39 - 1,608). The authors remarked that these estimates tended to reflect *N*_*S*_ rather than *N*_*e*_, which would be expected to be much larger than the former.

In this study, we show by computer simulations that the new software *currentNe2*, developed by Santiago et al. (2025), can solve this limitation of previous methods so that, with a sufficiently large sampling area, the metapopulation *N*_*e*_ can be estimated. Empirical data from maritime pine are used for illustration. Our results indicate that, incorporating samples from extended areas of its Western and Eastern European distribution, estimates of metapopulation *N*_*e*_ may reach values of up to tens of thousands of individuals, which contrasts with estimates of *N*_*e*_ assuming panmixia, generally below 1,000 individuals.

## Methods

### Simulation procedure

A custom C program was used to carry out the simulations. The model simulated was analogous to that considered by Neel et al. (2003) and Nunney (2016), consisting of a squared area with 300 × 300 cells in which there is a single individual at each of the cells. Thus, the population was composed of a total of 90,000 individuals. Following Neel et al. (2013), a breeding window (BW) is defined as the square area within which mating occurs at random. Thus, every generation, the individual of a given cell can be generated by two randomly chosen parents (excluding self-fertilization) from the breeding window from which the cell in question is central. The number of cells on one side of a breeding window took values of 5, 7, 9 and 15, so that the total breeding window would be BW = 25, 49, 81 and 225 cells. This breeding system was carried for 10,000 generations for up to 10 simulation replicates, and geometric mean estimates of *N*_*e*_ were obtained across replicates.

The genetic model consisted of a genome of 20 chromosomes with 1,020 potential loci each. In the initial generation, there was not variation at the loci, and this appeared by random recurrent mutation such that all loci were always segregating. Loci were biallelic (like single nucleotide polymorphism loci, SNP), and neutral, i.e. with no impact on the fitness of the individuals. A set of 680 multiallelic loci (34 per chromosome) at equal distances each, were also established in the last 15 generations at which a different allele was ascribed to each individual genome. This was aimed to estimate the identity-by-descent (IBD) homozygosity of each individual in the last 10 generations, and a measure of the IBD *N*_*e*_ obtained as *N*_*eIBD*_ = 10 / [2(*F*_*t*_ – *F*_*t*–10_) / (1 – *F*_*t*–10_)], where *F*_*t*_ is the average inbreeding coefficient of all individuals in the population at generation *t* = 10,000.

Recombination was set up at a uniform rate such that each chromosome was one Morgan long. Thus, simulations involved the processes of mutation, recombination, genetic drift and breeding system. At the last simulated generation, individuals were taken from a square area (the sampling area, SW), centred in the central cell of the whole population, with sides taking values of 5, 7, 9, 21, 41, 61, 81, 101, 151, 251 and 300, and therefore, SW = 25, 49, 441, 1,681, 3,721, 6,561, 10,201, 19,881, 32,761, 40401, 63,001 and 90,000, the latter being the whole population. All individuals were analysed in the sampling areas with less than 2,000 individuals, whereas a random sample of 2,000 individuals were taken from larger areas. Sampling was also made for the whole population but, instead of sampling individuals randomly across space, they were sampled considering 13 sampling points with size SW = 225 scattered uniformly in the whole area, as illustrated in Figure 1. In this case, also a random sample of 2,000 individuals was analysed, in order to be comparable with the fully random sampling scheme.

**Figure 1.**
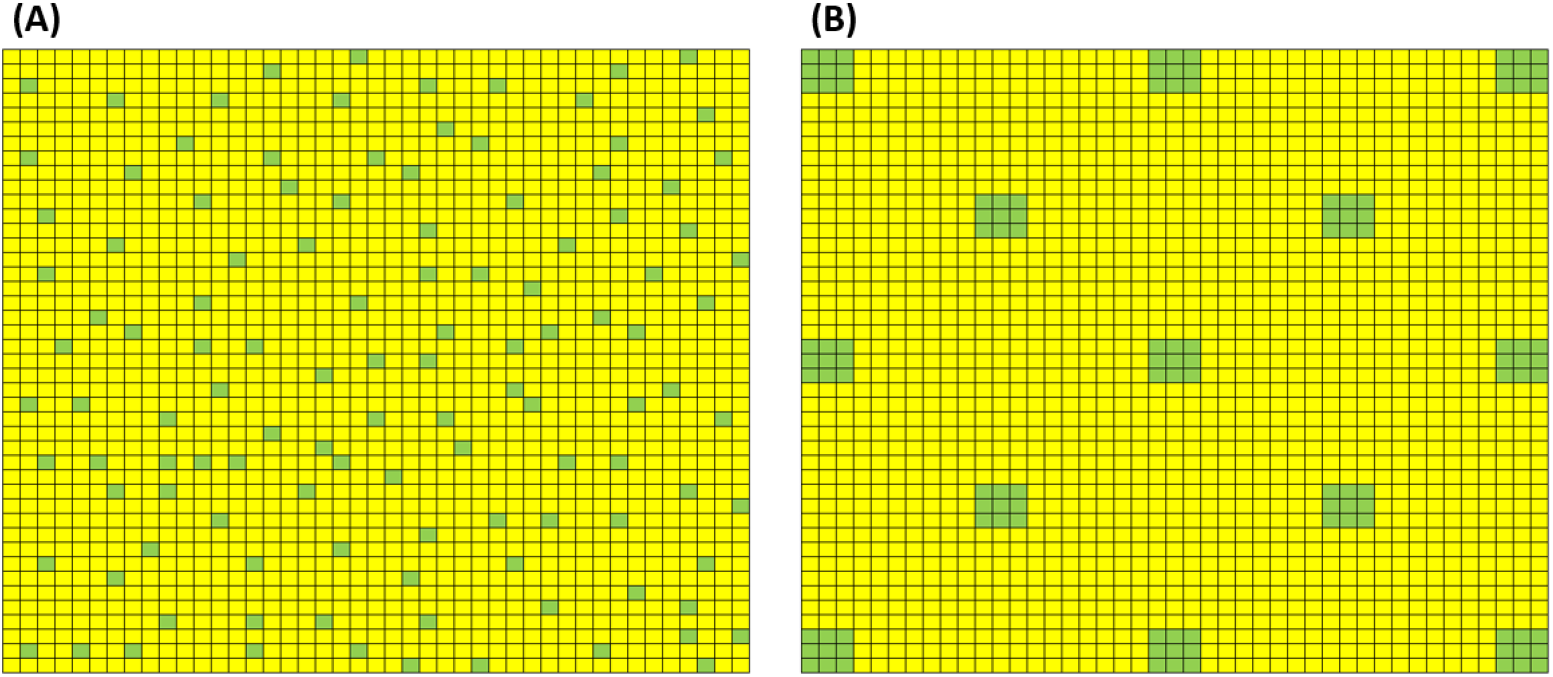
Sampling strategies followed in the simulations. (A) Sampled individuals are taken randomly from the whole population. (B) Sampled individuals are taken from 13 square areas (of size SW = 225) distributed across the whole population (the number of squares depicted is only for illustration purposes).

The neighbourhood size (*N*_*S*_) (Wright 1946, Rousset 2000) was obtained as *N*_*S*_ = 4πσ^2^*D*, where *D* is density (number of individuals per unit area) and σ is a measure of dispersal, given by the mean squared distance along one axis between birthplaces of parents and their offspring. The values of σ^2^ for BW = 25, 49, 81 and 225, were 2, 4, 6.7 and 18.7, respectively.

In order to evaluate the minimum and maximum values of *F*_*ST*_ across the population, this statistic was estimated as *F*_*ST*_ = (*H*_*T*_ – *H*_*S*_) / *H*_*T*_, where *H*_*S*_ is the expected heterozygosity within two breeding windows (BW) and *H*_*T*_ the expected heterozygosity assuming that both BWs are pooled. The minimum *F*_*ST*_ was obtained by considering two consecutive BWs, whereas the maximum *F*_*ST*_ was obtained by considering the two BWs in the opposite corners (upper right and lower left) of the full population area.

Individual genomic data obtained in the simulations was analysed with the software *currentNe2*, available at https://github.com/esrud/currentNe2, assuming panmixia (the default option) and population structure (the option –x). The software provides estimates of *N*_*e*_ from LD between pairs of SNPs across the entire genome, and their confidence limits obtained from a neural network approach. By assuming population structure, the software assumes an island model of migration and generates estimates of the metapopulation *N*_*e*_, the number of subpopulations (*s*), the migration rate (*m*) and the fixation index (*F*_*ST*_). The software has been shown to produce accurate estimates of these statistics for a variety of situations such as unequal subpopulation sizes, asymmetric migration, different sample sizes from subpopulations, etc. and applies also well for stepping stone models of migration (Santiago et al. 2025). Its performance has not been evaluated for continuously distributed populations with limited dispersal.

Estimates of *N*_*e*_ were also obtained by the software *LDNE* (Waples and Do 2008), which assumes panmixia in the population. These estimates assumed minor allele frequencies higher than 0.01 and were corrected for LD using the correction of Waples et al. (2016), which accounts for the number of chromosomes, by dividing the obtained estimate by 0.098 + 0.0219 ln(20).

### Empirical data of maritime pine

Genomic data (SNPs) from the European Western and Eastern range of maritime pine (*Pinus pinaster* Aiton, Pinaceae) (Milesi et al. 2024), and available at https://doi.org/10.57745/DV2X0M, were analysed with the software *currentNe2* to obtain estimates of contemporary *N*_*e*_ assuming panmixia or population structure. Sample geographical locations are shown in Figure 2 and Suppl. Table S1. A map of the sampling positions is also shown in Figure 1 of Milesi et al. (2024), as well as additional localities not used in this study. The original datasets (50% of maximum missing data) were filtered for a maximum of 5% missing data using the option -geno 0.05 with *PLINK v1*.*9* (Chang et al. 2015). Sampling in Milesi et al. (2024) was done in pairs of forest stands, to maximise local genetic diversity and sampled environmental variation. Pair samples in very close geographic proximity were analysed together, as assumed to be part of the same breeding population (see Figure 2 and Suppl. Table S1). The number of individuals and SNPs finally available for analysis are shown in Suppl. Table S2. Estimates of *F*_*ST*_ between localities were obtained with the software *vcftools* (Danacek et al. 2011). Estimates of *N*_*e*_ were obtained for each of the localities and for sets pooling localities as shown in Figure 2 for increasingly larger areas within Western and Eastern regions corresponding to two phylogeographic groups (Burban and Petit 2003).

**Figure 2.**
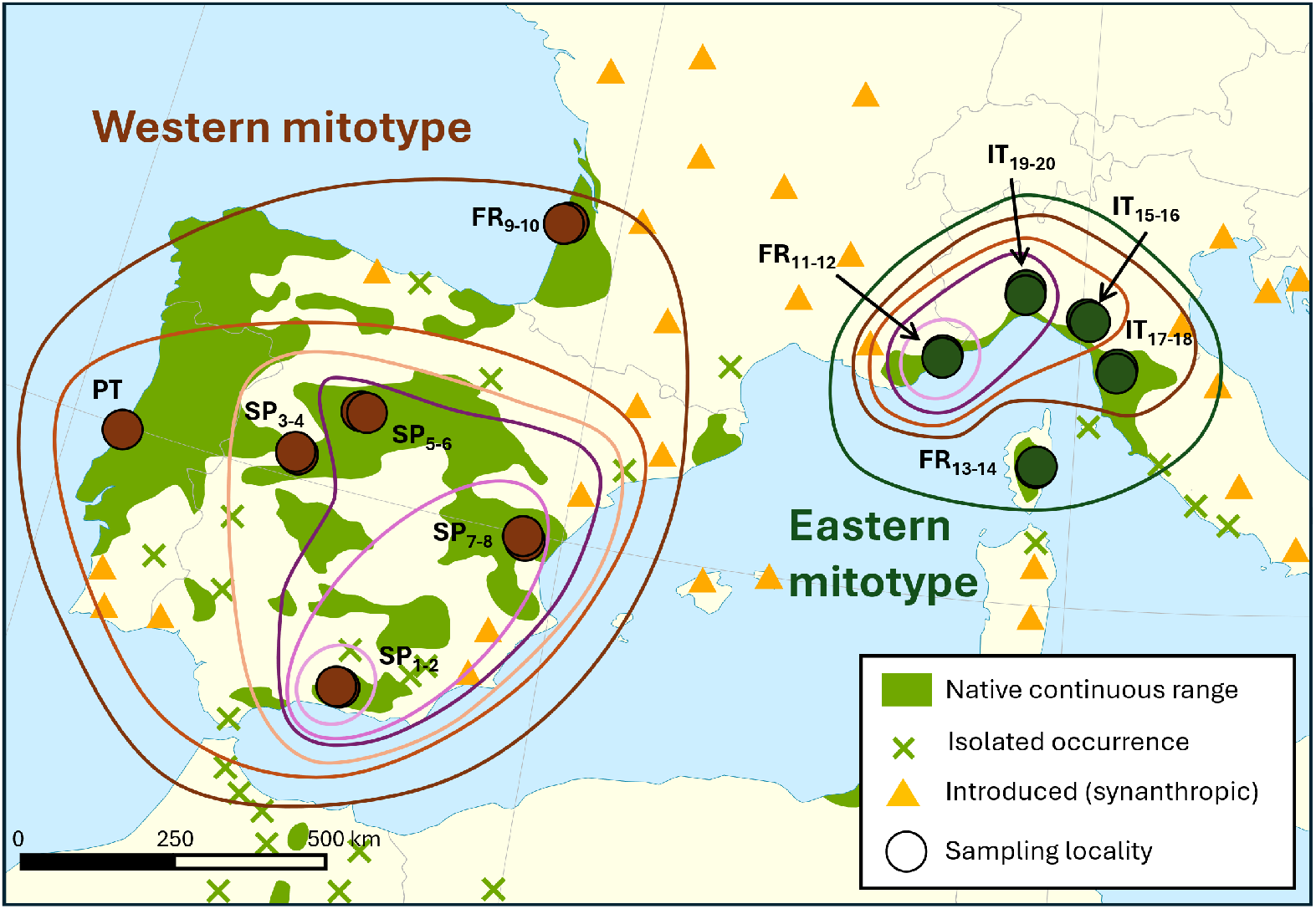
Map showing the sampling points. Subscript with two numbers indicate a sampling pair, i.e. two geographically close sampling locations analysed together (see Milesi et al. 2024 for details; see also Suppl. Table S1). Closed curves indicate the consecutive addition of sampled localities in extended Western and Eastern areas for *N*_*e*_ estimation (see also Suppl. Tables S5 and S6). The range of maritime pine is also shown (Caudullo et al. 2017).

## Results

### Simulation study

Average estimates of *N*_*e*_ obtained with the software *currentNe2* for four different breeding windows (BW) and a range of increasingly larger sampling windows (SW) are shown in Figure 3A. Standard errors of estimates are shown in Suppl. Table S3. The neighbourhood sizes corresponding to the four breeding windows were *Ns* = 25.13, 50.27, 83.78 and 546.67, respectively. Estimates assuming panmixia (red lines) started at values close to the neighbourhood size when SW was small, but increased as the SW increased towards an asymptotic value, higher for larger BW values. The estimates got slightly down for very large SW probably because of the edge effect of the cell grid (individuals in the edge have a number of potential parents lower than other individuals). All estimates were almost identical when obtained by the software *LDNE* (Waples and Do 2008) (Suppl. Figure S1). Estimates of *N*_*e*_ from the increase in the average inbreeding coefficient (IBD estimates) from simulated multilocus markers were around 270, 500, 800 and 2000 for BW of 25, 49, 81 and 225, respectively, irrespective of the sampling window size, which indicate the local *N*_*e*_ for each of the scenarios.

**Figure 3.**
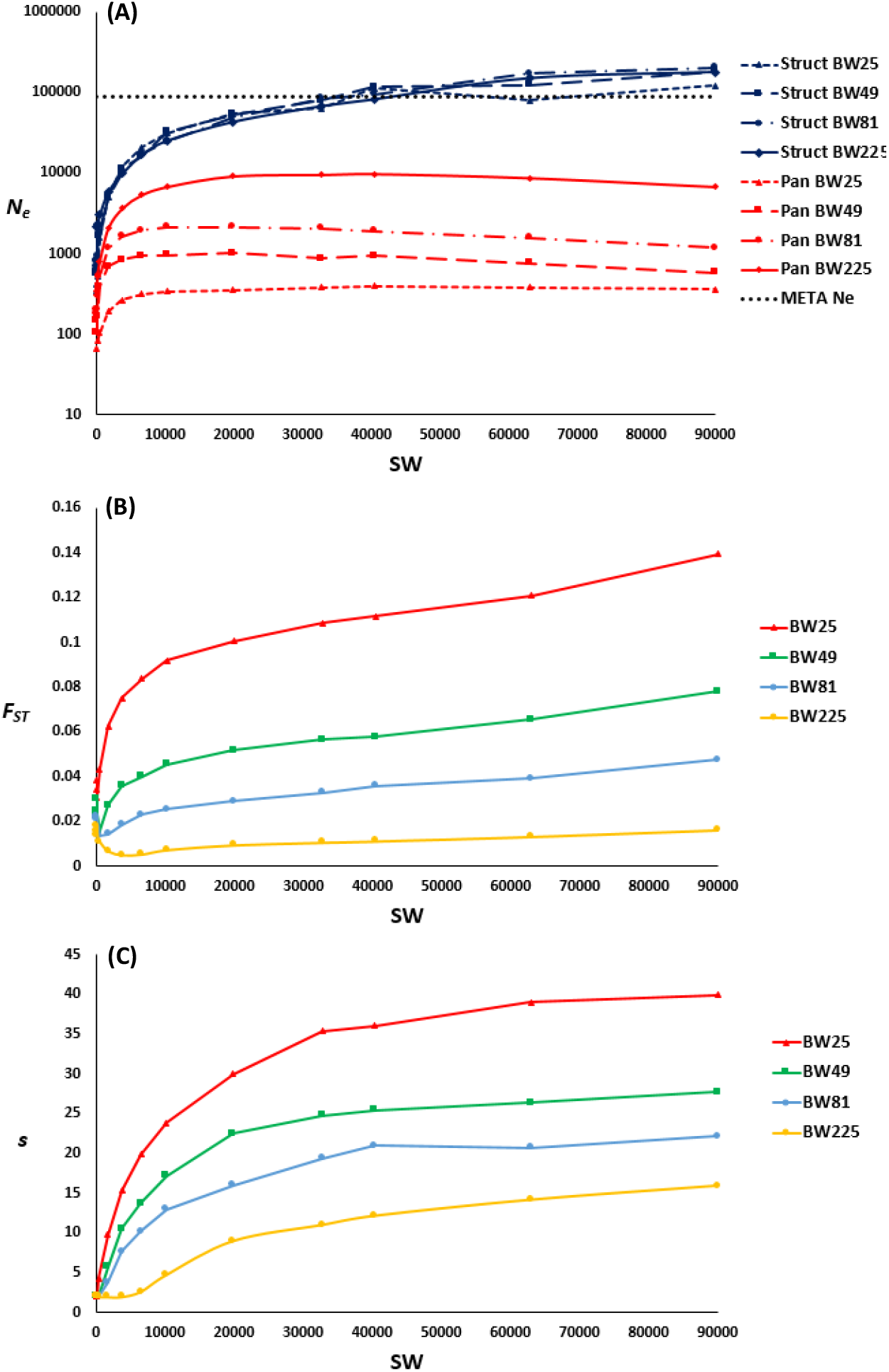
**(A)** Estimates of the effective population size *N*_*e*_ obtained with the software *currentNe2* (Santiago et al. 2025) for simulation results assuming panmixia (red lines) or population structure (blue lines) for different breeding windows (BW) and sampling windows (SW). **(B)** and **(C)** Estimates of the fixation index (*F*_*ST*_) and the number of subpopulations (*s*) obtained by the software.

Figure 3A also shows the estimates obtained by *currentNe2* when population structured is considered (blue lines). Again, estimates of *N*_*e*_ for small BW were close to the expected *Ns*, but the estimates increased towards the metapopulation *N*_*e*_ as the sample area was increased, and SW involving samples of individuals scattered across one third of the whole metapopulation distribution provided estimates of *N*_*e*_ reaching the total one. The estimates of *N*_*e*_ for very large SW were, however, overestimations (40% - 100%) of the total metapopulation size. The software gives estimates of the number of subpopulation (*s*) and the fixation index (*F*_*ST*_) assuming an island model. These are shown in Figure 3B-C, with increasing values of these statistics for increasing SW and BW. The estimates of *F*_*ST*_ from *currentNe2* when the whole population was sampled ranged between 0.016 (BW = 225) and 0.14 (BW = 25). These values are within the minimum and maximum values of *F*_*ST*_ obtained from the simulation data, *F*_*ST,min*_ = 0.003 and *F*_*ST,max*_ = 0.023 for BW= 225, and *F*_*ST,min*_ = 0.025 and *F*_*ST,max*_ = 0.204 for BW = 25.

Estimates of metapopulation *N*_*e*_ from *currentNe2* assuming population structure, obtained by concentrating the sampled individuals in 13 particular locations of SW = 225 individuals each, as shown in Figure 1B, provided values substantially lower than the whole population *N*_*e*_ (Table 1). For example, under the scenario with BW = 225 an estimated global *N*_*e*_ of 25,586 individuals was obtained, which is about one quarter the total number of individuals. For lower values of BW, estimates of metapopulation *N*_*e*_ were somewhat lower.

**Table 1.**
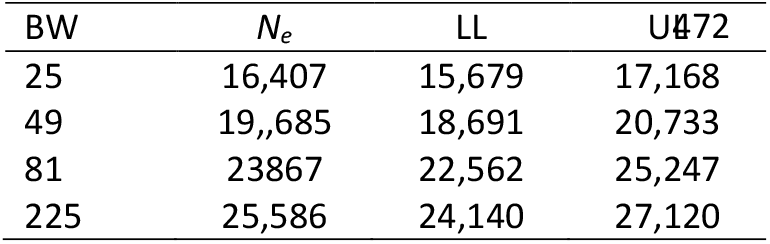
Average estimates (across up to ten replicates) of *N*_*e*_ obtained with the software *currentNe2* assuming population structure from sampling 13 specific points (SW = 225 individuals in each) within the whole population area, as shown in Figure 1B. Simulations assume different breeding windows (BW), and LL and UL denote lower and upper confidence limits of estimates of *N*_*e*_, respectively.

| BW | $N_e$ | LL | UL |
| --- | --- | --- | --- |
| 25 | 16,407 | 15,679 | 17,168 |
| 49 | 19,685 | 18,691 | 20,733 |
| 81 | 23,867 | 22,562 | 25,247 |
| 225 | 25,586 | 24,140 | 27,120 |

### Empirical application

Estimates of *N*_*e*_ were obtained with the software *currentNe2* for each of the samples shown in Figure 2. Estimates assuming panmixia had an average of 574 individuals (range 76 - 1,315) (Suppl. Table S4), while estimates assuming structure had an average of 5,161 (range 641 - 25,400), showing that the estimates considering structure were generally significantly larger than those assuming panmixia.

Estimates were then obtained by pooling samples from an increasing number of localities (as shown by the concentric circles in Figure 2 and Suppl. Table S5), following the expected dispersal of the species within its Western and Eastern distribution. The results show that *N*_*e*_ estimates assuming panmixia were generally low (average 195, range 38 – 811; Fig. 4A,B; Suppl. Table S6), whereas estimates were increasingly higher as more localities were included in the analysis (Fig. 4C,D; Suppl. Table S6), reaching values over 70,000 individuals in the Western area and over 120,000 individuals in the Eastern area. The estimates of *F*_*ST*_ ranged between 0.01 and 0.08, and the estimated number of subpopulations (*s*) was 6 for the Western range and 3 for the Eastern range (Suppl. Table S6), which agrees with the number of known gene pools in the species (Theraroz et al. 2024). The estimated *F*_*ST*_ by *currentNe2* for the whole set of Western populations was 0.073 and for the Eastern populations was 0.069. These values are very close to those obtained directly from the empirical data (0.066 for Western and 0.072 for Eastern regions; Suppl. Table S7), and those obtained previously by Milesi et al. (2024).

**Figure 4.**
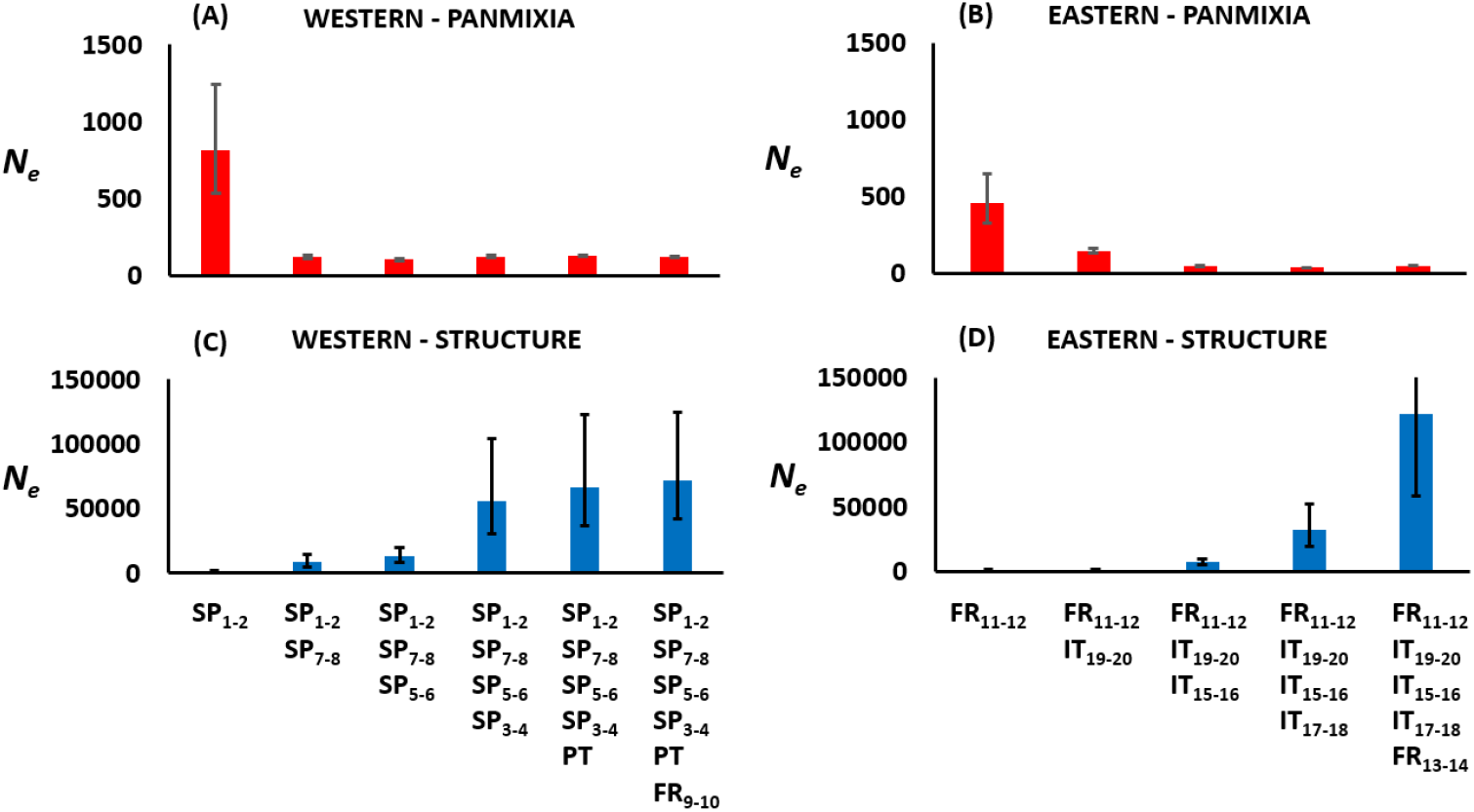
Estimates of effective population size (*N*_*e*_) and its lower and upper confidence 90% limits, obtained with the software *currentNe2* assuming panmixia (A, B, red bars) or assuming structure (C, D, blue bars) for empirical data of maritime pine populations. Estimates are obtained for sets of data for which localities are added consecutively considering the expected dispersal of the species. Western (A,C) and Eastern (B,D) maritime pine distribution localities in Europe, as shown in Fig. 2 and Suppl. Tables S5 and S6.

## Discussion

Our simulations considering panmixia agree with analogous ones obtained by Neel et al. (2013) from LD estimates and by Nunney (2016) using the temporal method of estimation of *N*_*e*_, based on the change in allele frequency of markers over generations. Neither the methods above nor *currentNe2* assuming panmixia can estimate the global metapopulation *N*_*e*_ (in this case 90,000 individuals), even if the whole population area were sampled. Rather, the asymptotic *N*_*e*_ assuming panmixia (281, 580, 1,169 and 6,725 for BW = 25, 49, 81 and 225, respectively) were of the order of the local *N*_*e*_ estimates obtained from the increase in the average inbreeding (IBD) obtained from multilocus markers (around 270, 500, 800 and 2,000 for BW = 25, 49, 81 and 225, respectively). This is a consequence of the estimation of *N*_*e*_ in structured populations by methods assuming panmixia (Waples and England 2011, Ryman et al. 2009, Novo et al. 2023, Kardos and Waples 2024), because structure induces additional LD, even between sites on different chromosomes. If this effect is ignored, *N*_*e*_ will be significantly underestimated. This explains the very low ratios *N*_*e*_/*N* found in large continuously distributed populations, such as those for marine fish and forest tree species (Neel et al. 2013, Marandel et al. 2019, Waples 2024, Fedorca et al. 2024, Gargiulo et al. 2024, Santos-del-Blanco et al. 2022).

Neel et al. (2013) assumed also a 300 × 300 cell grid scheme for a population of *N* = 90,000 individuals run for 2,000 generations. They simulated 100 multiallelic loci initially with 99 alleles (analogous to microsatellites), and estimated *N*_*e*_ (*N*_*b*_ in their notation) for samples of 100 individuals in the last 100 generations and for 100 replicates using the software *LDNE* (Waples and Do 2008). To avoid edge effects, they only considered centred sampling windows with side up to 66 cells, which implies SW = 4,356. The estimated *N*_*e*_ for this largest SW and BW = 25, 49 and 81 reached values <200, <400 and <800, respectively. In our simulations, we run a similar scenario for 10,000 generations, considering markers analogous to SNPs and analysing only the last generation for up to ten replicates with a maximal sample size of 2,000 individuals. Our estimates of *N*_*e*_ with *currentNe2* assuming panmixia and those obtained with *LDNE* were almost identical to each other, although the latter were a bit lower than the former (Suppl. Fig. S1), probably because Neel et al. (2013) presented the harmonic mean of estimates from many replicates (100 ×100), while we used the geometric mean of ten replicates.

We considered SW from a small one up to the whole population area. The overall results from Neel et al. (2013) and those of the present study assuming panmixia are fully concordant in that spatial variation in a continuous population can severely bias estimates of *N*_*e*_ from LD, with estimates of *N*_*e*_ only a few times larger than the neighbourhood size *Ns*, but much lower than the metapopulation *N*_*e*_. The results are also concordant with those of Nunney (2016), who considered a similar breeding model with lower population sizes (up to 4,096 individuals) and estimated *N*_*e*_ from the change in allele frequency of markers in a 32-generation period. For example, for the scenario with a neighbourhood size of *Ns* = 64, the estimated *N*_*e*_ (201) was larger than *Ns*, but substantially lower than *N*_*e*_ for the entire population (1,365).

The estimates of *N*_*e*_ with the software *currentNe2* assuming population structure are expected to provide estimates of *N*_*e*_ lower or non-significantly different from those assuming panmixia when the population is a closed one or there is subdivision but the migration rate is very large between subpopulations, with *N*_*e,spop*_*m* > 10, where *N*_*e,spop*_ is the subpopulation size and *m* the migration rate (Santiago et al. 2025). If the population is structured with *N*_*e,spop*_*m* < 10 the estimates assuming structure should be higher than those assuming panmixia, and yield estimates of the metapopulation *N*_*e*_. From Figure 3 it can be seen that estimates close to the total population size (*N* = 90,000) were obtained when the sampling area considered was about one third of the total area for the population. In the limiting case of sampling over the whole population area, estimates were substantially larger than *N* = 90,000, what denotes an overestimation of *N*_*e*_. In contrast, when the estimates of *N*_*e*_ were obtained from particular sampling points, even if scattered across the whole population area (Figure 1B), the global estimates obtained were an underestimation of the metapopulation *N*_*e*_ (Table 1). These results remark the importance of obtaining appropriate samples specifically devoted for the estimation of *N*_*e*_ across the distribution area of the species, rather than trying to reuse samples obtained for other purposes.

Estimates of the putative number of subpopulations (in an Island model), the migration rate and the fixation index (*F*_*ST*_) were also obtained by the software. In our analyses, the number of subpopulations and *F*_*ST*_ were increased up to a limiting value of *s* ≈ 40 and *F*_*ST*_ ≈ 0.14 in the scenario of BW = 25 (Figure 3B,C). Importantly, as suggested by Wright (1943), and shown by Nunney (2016), a neighbourhood in a continuous distributed population would be analogous to an island of the island model if neighbourhoods were sampled at random. In fact, the estimates of *F*_*ST*_ obtained by *currentNe2* were within the limits of the minimum and maximum estimates obtained between the closest and the most distant breeding windows.

The application of the method to empirical data from European samples of maritime pine shows that estimates assuming panmixia and including increasingly larger areas within the distribution of the species, produced estimates of *N*_*e*_ rather low, of the order of a few hundred at most, in agreement with previous studies (Santos-del-Blanco et al. 2022). In contrast, when the method assumed population structure, estimates pooling samples from wide geographic areas provided estimates of *N*_*e*_ over 70,000 (Western distribution) and over 120,000 (Eastern distribution) individuals, more coherent with populations of very large census sizes. Estimates of this order are in agreement with alternative estimates obtained from the site frequency spectrum of allele markers (Jaramillo-Correa et al. 2020), although this method is not very appropriate for estimating contemporary *N*_*e*_ (Nadachowska-Brzyska et al. 2022).

The estimated neighbourhood size in maritime pine is around 100-150 individuals (de-Lucas et al. 2008, 2009), which would be comparable with our simulation results for BW = 81-225. The values of *F*_*ST*_ between pine localities are of the order of 0.07 (Suppl. Table S7), and these are also similar to the values obtained in the simulations (Figure 3B), so that a comparison between simulation results and empirical ones looks appropriate. The simulation results show that, when sampling is sufficiently large across the whole population area, the estimate of metapopulation *N*_*e*_ can be an overestimate of the true *N*_*e*_. This may suggest that the estimates of metapopulation *N*_*e*_ in maritime pine can also be overestimations. However, the simulations also show that, when sampling is made in a few particular localities, even if well scattered across the population area (Figure 1B), the estimates of metapopulation *N*_*e*_ are underestimations of the global *N*_*e*_ (Table 1). Sampling in the empirical data was carried out in particular localities (Figure 2), and this would point out towards an underestimation of metapopulation *N*_*e*_ of pines. Despite these uncertainties about possible over or underestimations, it is clear that the metapopulation *N*_*e*_ of pines is of the order of tens of thousands individuals rather than below 1,000, as previously obtained with methods which assume panmixia. Nevertheless, even these large estimates of metapopulation *N*_*e*_ suggest that the ratio *N*_*e*_ / *N* in this species is small, as the number of individuals is, in rough numbers, on the order of 5 billion trees, assuming an average density of 1,200 trees/ha and the current over 4.2 million hectares of maritime pine forests in Europe.

Our results are very relevant from a conservation point of view, because a consensus rule is that populations with *N*_*e*_ values lower than 500-1,000 can be considered under the risk of extinction (Franklin 1980, Frankham et al. 2014, Pérez-Pereira et al. 2022). This has been recently translated into policy by establishing *N*_*e*_ > 500 as a headline indicator of the Kunming-Montreal Global Biodiversity Framework under the UN’s Convention on Biological Diversity (2022). Thus, accounting for population structure and migration may have, therefore, a profound impact on conservation decisions.

## Acknowledgements

This study forms part of the Marine Science Programme (ThinkInAzul) supported by the Ministerio de Ciencia e Innovación and Xunta de Galicia with funding from the European Union NextGenerationEU (PRTRC17.I1) and European Maritime and Fisheries Fund. We also acknowledge support from grants PID2020-114426GB-C21 (funded by MCIN/AEI/10.13039/501100011033), Xunta de Galicia (ED431C 2024/22), Centro Singular de Investigación de Galicia accreditation 2024-2027 (ED431G 2023/07), “ERDF A way of making Europe” and the European Union’s Horizon 2020 research and innovation programme under grant agreement No. 862221 (FORGENIUS). Views and opinions expressed are, however, those of the authors only and do not necessarily reflect those of the European Union. Neither the European Union nor the granting authority can be held responsible for them. Part of the analyses of this project were run using computing resources and technical support provided by CESGA (Centro de Supercomputación de Galicia). Funding for open access charge: Universidade de Vigo/CISUG. Open Access funding provided thanks to the CRUE-CSIC agreement with Wiley.

## Authors contribution statement

A.C. conceived the investigation and carried out the simulations. SGM provided the empirical data. All co-authors contributed to the analysis of data, interpretation of results and writing of the manuscript.

## Artificial Intelligence

AI has not been used in any way to carry out the study or to write the paper.

## Data availability

The datasets used for the empirical analysis are available at https://doi.org/10.57745/DV2X0M.

## Code availability

The C program code and the output files of analyses are available at GitHub address: https://github.com/armando-caballero/Ne-for-continuously-distributed-populations

## SUPPLEMENTARY INFORMATION

**Supplementary Table S1.** Coordinates of samples from maritime pine (*Pinus pinaster* Aiton) obtained by Milesi et al. (2024) and analysed in this study. Sample pairs in close geographical proximity were combined for analysis (i.e., SP_1-2_ involves samples SP_01 and SP_02; and so on).

| Country | ID | Latitude | Longitude | Sample pair or locality |
| --- | --- | --- | --- | --- |
| Spain | SP_01 | 36.8274076 | -3.9408404 | SP<sub>1-2</sub> |
|  | SP_02 | 36.834906 | -3.9239528 |  |
|  | SP_03 | 40.2451584 | -5.1216644 | SP<sub>3-4</sub> |
|  | SP_04 | 40.1888112 | -5.1082268 |  |
|  | SP_05 | 41.3361088 | -4.2459272 | SP<sub>5-6</sub> |
|  | SP_06 | 41.3412568 | -4.2351404 |  |
|  | SP_07 | 39.9172976 | -0.394194 | SP<sub>7-8</sub> |
|  | SP_08 | 39.912428 | -0.38906 |  |
| Portugal | PT | 39.783333 | -8.9575 | PT |
| France | FR_09 | 44.9685918 | -1.1639783 | FR<sub>9-10</sub> |
|  | FR_10 | 44.77984964 | -1.22913708 |  |
|  | FR_11 | 43.23320728 | 6.36536648 | FR<sub>11-12</sub> |
|  | FR_12 | 43.21737628 | 6.3600182 |  |
|  | FR_13 | 41.754864 | 9.2118156 | FR<sub>13-14</sub> |
|  | FR_14 | 41.8162124 | 9.2603792 |  |
| Italy | IT_15 | 44.0512058 | 9.970569 | IT<sub>15-16</sub> |
|  | IT_16 | 44.01190268 | 10.18390916 |  |
|  | IT_17 | 43.1316902 | 11.24342688 | IT<sub>17-18</sub> |
|  | IT_18 | 43.1391764 | 11.3489856 |  |
|  | IT_19 | 44.41752003 | 8.671061 | IT<sub>19-20</sub> |
|  | IT_20 | 44.55097224 | 8.64537032 |  |

**Supplementary Table S2.**
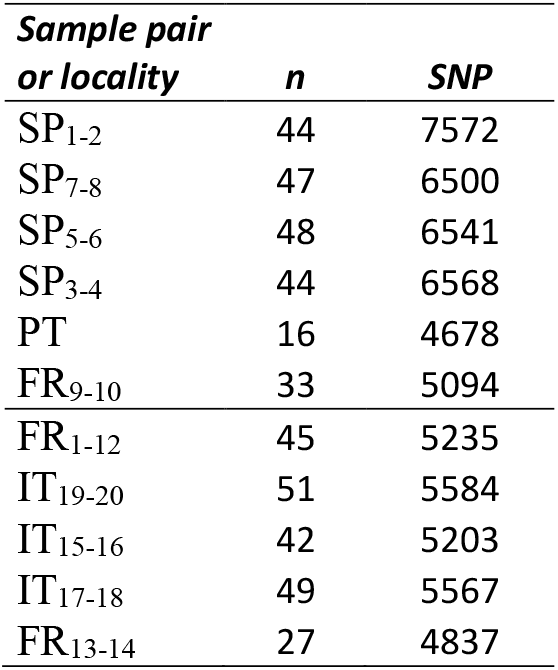
Number of sampled individuals (*n*) and number of SNPs used in the analysis of maritime pine populations.

**Supplementary Table S3.** Estimates of effective population size (*N*_*e*_) obtained with the software *currentNe2* (Santiago et al. 2025) assuming panmixia or assuming population structure and their upper (UL) and lower (LL) 90% confidence limits, for different breeding windows (BW) and sampling windows (SW).

| BW | SW | Panmixia |  |  | Structure |  |  |
| --- | --- | --- | --- | --- | --- | --- | --- |
| | | $N_e$ | LL | UL | $N_e$ | LL | UL |
| 25 | 25 | **67** | 49 | 92 | **412** | 216 | 787 |
| 25 | 49 | **83** | 68 | 100 | **337** | 243 | 467 |
| 25 | 81 | **104** | 91 | 118 | **522** | 407 | 670 |
| 25 | 441 | **192** | 186 | 197 | **1571** | 1446 | 1707 |
| 25 | 1681 | **265** | 265 | 265 | **4940** | 4777 | 5108 |
| 25 | 3721 | **313** | 313 | 313 | **11772** | 11274 | 12291 |
| 25 | 6561 | **342** | 342 | 342 | **20901** | 19754 | 22116 |
| 25 | 10201 | **352** | 352 | 352 | **30964** | 28938 | 33133 |
| 25 | 19881 | **378** | 378 | 378 | **54630** | 50006 | 59682 |
| 25 | 32761 | **395** | 395 | 395 | **63355** | 57954 | 69260 |
| 25 | 40401 | **385** | 385 | 385 | **111327** | 98739 | 125521 |
| 25 | 63001 | **360** | 360 | 360 | **79758** | 72375 | 87895 |
| 25 | 90000 | **281** | 281 | 281 | **122484** | 108556 | 138199 |
| 49 | 25 | **105** | 73 | 152 | **574** | 282 | 1168 |
| 49 | 49 | **145** | 115 | 183 | **581** | 391 | 863 |
| 49 | 81 | **165** | 141 | 192 | **571** | 444 | 736 |
| 49 | 441 | **392** | 375 | 409 | **1638** | 1507 | 1780 |
| 49 | 1681 | **684** | 684 | 684 | **5002** | 4840 | 5170 |
| 49 | 3721 | **838** | 838 | 838 | **11192** | 10738 | 11666 |
| 49 | 6561 | **936** | 936 | 936 | **16244** | 15462 | 17067 |
| 49 | 10201 | **954** | 954 | 954 | **32042** | 29957 | 34271 |
| 49 | 19881 | **1005** | 1005 | 1005 | **53606** | 49324 | 58259 |
| 49 | 32761 | **870** | 870 | 870 | **78990** | 71644 | 87090 |
| 49 | 40401 | **931** | 931 | 931 | **117073** | 104369 | 131324 |
| 49 | 63001 | **763** | 763 | 763 | **124273** | 110538 | 139714 |
| 49 | 90000 | **580** | 580 | 580 | **201078** | 173414 | 233155 |
| 81 | 25 | **181** | 115 | 285 | **694** | 325 | 1482 |
| 81 | 49 | **204** | 157 | 265 | **782** | 503 | 1217 |
| 81 | 81 | **202** | 171 | 239 | **918** | 679 | 1242 |
| 81 | 441 | **532** | 506 | 559 | **1945** | 1781 | 2124 |
| 81 | 1681 | **1159** | 1159 | 1159 | **4938** | 4781 | 5099 |
| 81 | 3721 | **1626** | 1626 | 1626 | **9856** | 9487 | 10238 |
| 81 | 6561 | **1923** | 1923 | 1923 | **17148** | 16307 | 18033 |
| 81 | 10201 | **2122** | 2120 | 2124 | **24879** | 23438 | 26408 |
| 81 | 19881 | **2124** | 2124 | 2124 | **48234** | 44559 | 52211 |
| 81 | 32761 | **2073** | 2073 | 2073 | **84210** | 76183 | 93084 |
| 81 | 40401 | **1919** | 1919 | 1919 | **94806** | 85419 | 105226 |
| 81 | 63001 | **1586** | 1586 | 1586 | **171360** | 149765 | 196069 |
| 81 | 90000 | **1169** | 1169 | 1169 | **203225** | 176160 | 234449 |

**Supplementary Table S3.** Continuation.
|  |  |  |  |  |  |  |  |
| --- | --- | --- | --- | --- | --- | --- | --- |
| 225 | 25 | **348** | 196 | 618 | **858** | 379 | 1941 |
| 225 | 49 | **304** | 225 | 409 | **2168** | 1143 | 4114 |
| 225 | 81 | **528** | 416 | 669 | **2358** | 1536 | 3620 |
| 225 | 441 | **806** | 760 | 855 | **3065** | 2761 | 3401 |
| 225 | 1681 | **2055** | 2027 | 2083 | **5887** | 5686 | 6095 |
| 225 | 3721 | **3616** | 3555 | 3679 | **9879** | 9518 | 10254 |
| 225 | 6561 | **5321** | 5192 | 5454 | **16789** | 15987 | 17630 |
| 225 | 10201 | **6701** | 6510 | 6898 | **25169** | 23726 | 26700 |
| 225 | 19881 | **9121** | 8808 | 9444 | **43324** | 40204 | 46687 |
| 225 | 32761 | **9501** | 9169 | 9844 | **67663** | 61823 | 74055 |
| 225 | 40401 | **9753** | 9409 | 10110 | **82408** | 74732 | 90874 |
| 225 | 63001 | **8672** | 8387 | 8967 | **150429** | 132741 | 170473 |
| 225 | 90000 | **6725** | 6537 | 6918 | **179255** | 156618 | 205164 |

**Supplementary Table S4.** Estimates of effective population size (*N*_*e*_) and its lower (LL) and upper (UL) confidence 90% limits, obtained with the software *currentNe2* assuming panmixia or assuming structure (option -x) for empirical data of maritime pine samples. *F*_*ST*_: estimate of Wright’s (1943) fixation index; *m*: estimate of migration rate; *s*: estimate of the number of subpopulations assuming an island model. Scenarios for which the interval coefficients of *N*_*e*_ assuming structure overlap with those assuming panmixia are considered non-structured populations (Santiago et al. 2025) (*s* = 1).

| *Sample pair or locality* | PANMIXIA |  |  | STRUCTURE |  |  |  |  |  |
| --- | --- | --- | --- | --- | --- | --- | --- | --- | --- |
| | $N_e$ | LL | UL | $F_{ST}$ | $m$ | $s$ | $N_e$ | LL | UL |
| SP<sub>1-2</sub> | 811 | 530 | 1242 | 0.0265 | 0.2137 | 1 | 1459 | 860 | 2475 |
| SP<sub>7-8</sub> | 306 | 231 | 404 | 0.0413 | 0.0166 | 6 | 1524 | 911 | 2550 |
| SP<sub>5-6</sub> | 932 | 614 | 1414 | * |  |  |  |  |  |
| SP<sub>3-4</sub> | 959 | 607 | 1515 | 0.0158 | 0.0069 | 13 | 25400 | 5410 | 119247 |
| PT | 1315 | 398 | 4347 | 0.0333 | 0.0019 | 1 | 12555 | 1014 | 155421 |
| FR<sub>9-10</sub> | 699 | 420 | 1163 | 0.0001 | 0.4486 | 19 | 5554 | 1876 | 16444 |
| FR<sub>1-12</sub> | 461 | 329 | 647 | 0.0125 | 0.4499 | 1 | 1020 | 645 | 1613 |
| IT<sub>19-20</sub> | 97 | 83 | 114 | 0.0300 | 0.0065 | 2 | 641 | 457 | 899 |
| IT<sub>15-16</sub> | 282 | 206 | 385 | 0.0398 | 0.0528 | 12 | 1200 | 699 | 2059 |
| IT<sub>17-18</sub> | 76 | 65 | 88 | 0.0396 | 0.0020 | 2 | 1549 | 947 | 2533 |
| FR<sub>13-14</sub> | 373 | 232 | 600 | 0.0071 | 0.4498 | 1 | 711 | 387 | 1308 |

\* $N_e$ could not be estimated because the LD was smaller for linked than for unlinked markers.

**Supplementary Table S5.**
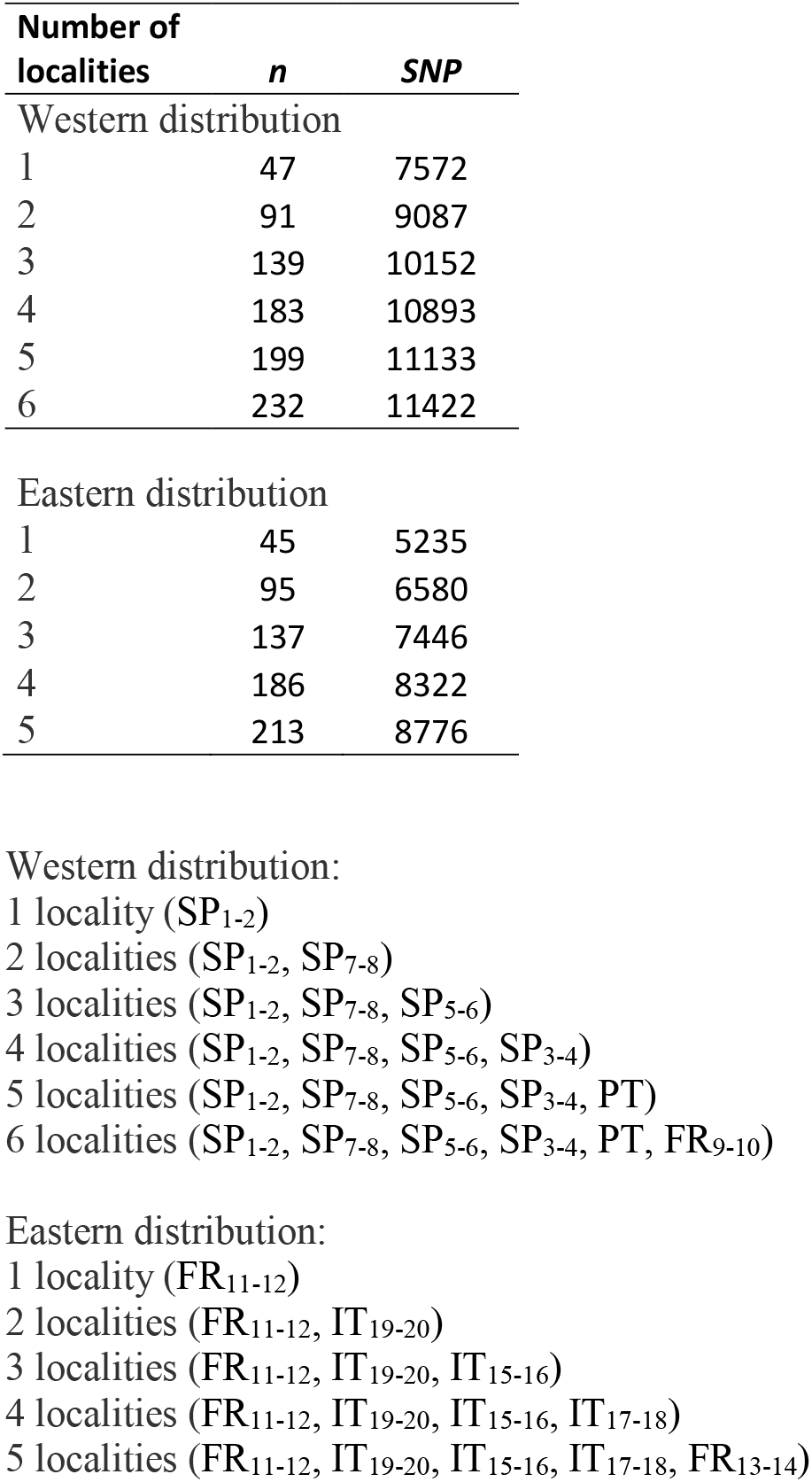
Number of individuals (*n*) and number of SNPs used in the analysis of maritime pine populations pooling localities.

| Number of localities | *n* | *SNP* |
| --- | --- | --- |
| Western distribution |  |  |
| 1 | 47 | 7572 |
| 2 | 91 | 9087 |
| 3 | 139 | 10152 |
| 4 | 183 | 10893 |
| 5 | 199 | 11133 |
| 6 | 232 | 11422 |
| Eastern distribution |  |  |
| 1 | 45 | 5235 |
| 2 | 95 | 6580 |
| 3 | 137 | 7446 |
| 4 | 186 | 8322 |
| 5 | 213 | 8776 |

Western distribution: 1 locality (SP<sub>1-2</sub>) 2 localities (SP<sub>1-2</sub>, SP<sub>7-8</sub>) 3 localities (SP<sub>1-2</sub>, SP<sub>7-8</sub>, SP<sub>5-6</sub>) 4 localities (SP<sub>1-2</sub>, SP<sub>7-8</sub>, SP<sub>5-6</sub>, SP<sub>3-4</sub>) 5 localities (SP<sub>1-2</sub>, SP<sub>7-8</sub>, SP<sub>5-6</sub>, SP<sub>3-4</sub>, PT) 6 localities (SP<sub>1-2</sub>, SP<sub>7-8</sub>, SP<sub>5-6</sub>, SP<sub>3-4</sub>, PT, FR<sub>9-10</sub>) Eastern distribution: 1 locality (FR<sub>11-12</sub>) 2 localities (FR<sub>11-12</sub>, IT<sub>19-20</sub>) 3 localities (FR<sub>11-12</sub>, IT<sub>19-20</sub>, IT<sub>15-16</sub>) 4 localities (FR<sub>11-12</sub>, IT<sub>19-20</sub>, IT<sub>15-16</sub>, IT<sub>17-18</sub>) 5 localities (FR<sub>11-12</sub>, IT<sub>19-20</sub>, IT<sub>15-16</sub>, IT<sub>17-18</sub>, FR<sub>13-14</sub>)

**Supplementary Table S6.** Estimates of effective population size (*N*_*e*_) and its lower (LL) and upper (UL) confidence 90% limits, obtained with the software *currentNe2* assuming panmixia or assuming structure for empirical data of maritime pine samples pooling data from different localities. *F*_*ST*_: estimate of Wright’s (1943) fixation index; *m*: estimate of migration rate; *s*: estimate of the number of subpopulations assuming and island model. Scenarios for which the interval coefficients of *N*_*e*_ assuming structure overlap with those assuming panmixia are considered non-structured populations (Santiago et al. 2025) (*s* = 1).

| PANMIXIA |  |  |  | STRUCTURE |  |  |  |  |  |
| --- | --- | --- | --- | --- | --- | --- | --- | --- | --- |
| Number of localities | $N_e$ | LL | UL | $F_{ST}$ | $m$ | $s$ | $N_e$ | LL | UL |
| Western distribution |  |  |  |  |  |  |  |  |  |
| 1 | 811 | 530 | 1242 | 0.0265 | 0.2137 | 1 | 1459 | 860 | 2475 |
| 2 | 119 | 108 | 133 | 0.0644 | 0.0015 | 5 | 8382 | 4856 | 14471 |
| 3 | 101 | 95 | 108 | 0.0698 | 0.0009 | 5 | 12922 | 8293 | 20135 |
| 4 | 123 | 117 | 130 | 0.0631 | 0.0002 | 5 | 56217 | 30330 | 104198 |
| 5 | 127 | 121 | 134 | 0.0690 | 0.0002 | 6 | 66819 | 36295 | 123013 |
| 6 | 117 | 112 | 121 | 0.0728 | 0.0002 | 6 | 72008 | 41661 | 124462 |
| Eastern distribution |  |  |  |  |  |  |  |  |  |
| 1 | 461 | 329 | 647 | 0.0125 | 0.4499 | 1 | 1020 | 645 | 1613 |
| 2 | 145 | 130 | 162 | 0.0342 | 0.0079 | 3 | 1238 | 963 | 1591 |
| 3 | 48 | 46 | 50 | 0.0695 | 0.0007 | 3 | 6888 | 4822 | 9838 |
| 4 | 38 | 37 | 39 | 0.0787 | 0.0001 | 3 | 32044 | 19602 | 52384 |
| 5 | 49 | 48 | 50 | 0.0693 | 0.0000 | 3 | 121840 | 58732 | 252759 |

Western distribution: 1 locality (SP<sub>1-2</sub>) 2 localities (SP<sub>1-2</sub>, SP<sub>7-8</sub>) 3 localities (SP<sub>1-2</sub>, SP<sub>7-8</sub>, SP<sub>5-6</sub>) 4 localities (SP<sub>1-2</sub>, SP<sub>7-8</sub>, SP<sub>5-6</sub>, SP<sub>3-4</sub>) 5 localities (SP<sub>1-2</sub>, SP<sub>7-8</sub>, SP<sub>5-6</sub>, SP<sub>3-4</sub>, PT) 6 localities (SP<sub>1-2</sub>, SP<sub>7-8</sub>, SP<sub>5-6</sub>, SP<sub>3-4</sub>, PT, FR<sub>9-10</sub>)
Eastern distribution: 1 locality (FR<sub>11-12</sub>) 2 localities (FR<sub>11-12</sub>, IT<sub>19-20</sub>) 3 localities (FR<sub>11-12</sub>, IT<sub>19-20</sub>, IT<sub>15-16</sub>) 4 localities (FR<sub>11-12</sub>, IT<sub>19-20</sub>, IT<sub>15-16</sub>, IT<sub>17-18</sub>) 5 localities (FR<sub>11-12</sub>, IT<sub>19-20</sub>, IT<sub>15-16</sub>, IT<sub>17-18</sub>, FR<sub>13-14</sub>)

**Supplementary Table S7.**
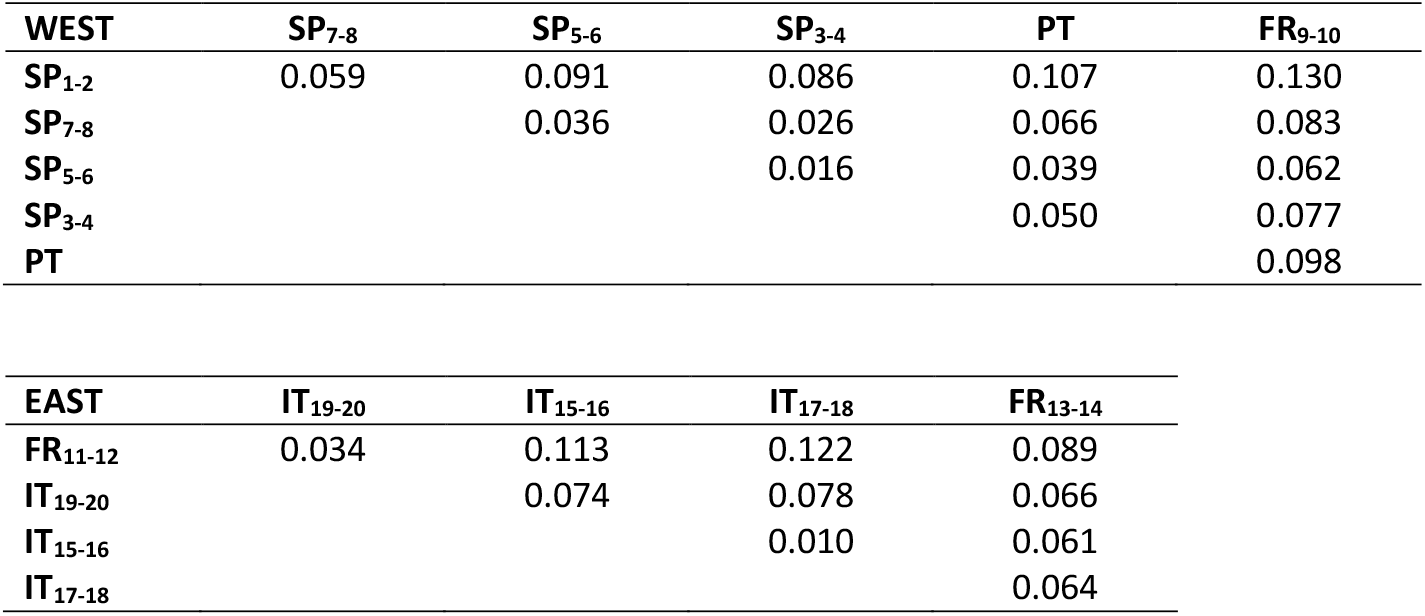
Estimates of *F*_*ST*_ between pairs of European maritime pine populations obtained by the software *vcftools* (Danacek et al. 2011). Symbols of populations as shown in Figure 2 and Supp. Tables S1-S2. Estimates considering all populations were 0.066 for the Western localities and 0.072 for the Eastern ones

| **WEST** | **SP<sub>7-8</sub>** | **SP<sub>5-6</sub>** | **SP<sub>3-4</sub>** | **PT** | **FR<sub>9-10</sub>** |
| --- | --- | --- | --- | --- | --- |
| **SP<sub>1-2</sub>** | 0.059 | 0.091 | 0.086 | 0.107 | 0.130 |
| **SP<sub>7-8</sub>** |  | 0.036 | 0.026 | 0.066 | 0.083 |
| **SP<sub>5-6</sub>** |  |  | 0.016 | 0.039 | 0.062 |
| **SP<sub>3-4</sub>** |  |  |  | 0.050 | 0.077 |
| **PT** |  |  |  |  | 0.098 |

| **EAST** | **IT<sub>19-20</sub>** | **IT<sub>15-16</sub>** | **IT<sub>17-18</sub>** | **FR<sub>13-14</sub>** |
| --- | --- | --- | --- | --- |
| **FR<sub>11-12</sub>** | 0.034 | 0.113 | 0.122 | 0.089 |
| **IT<sub>19-20</sub>** |  | 0.074 | 0.078 | 0.066 |
| **IT<sub>15-16</sub>** |  |  | 0.010 | 0.061 |
| **IT<sub>17-18</sub>** |  |  |  | 0.064 |

**Supplementary Figure S1.**
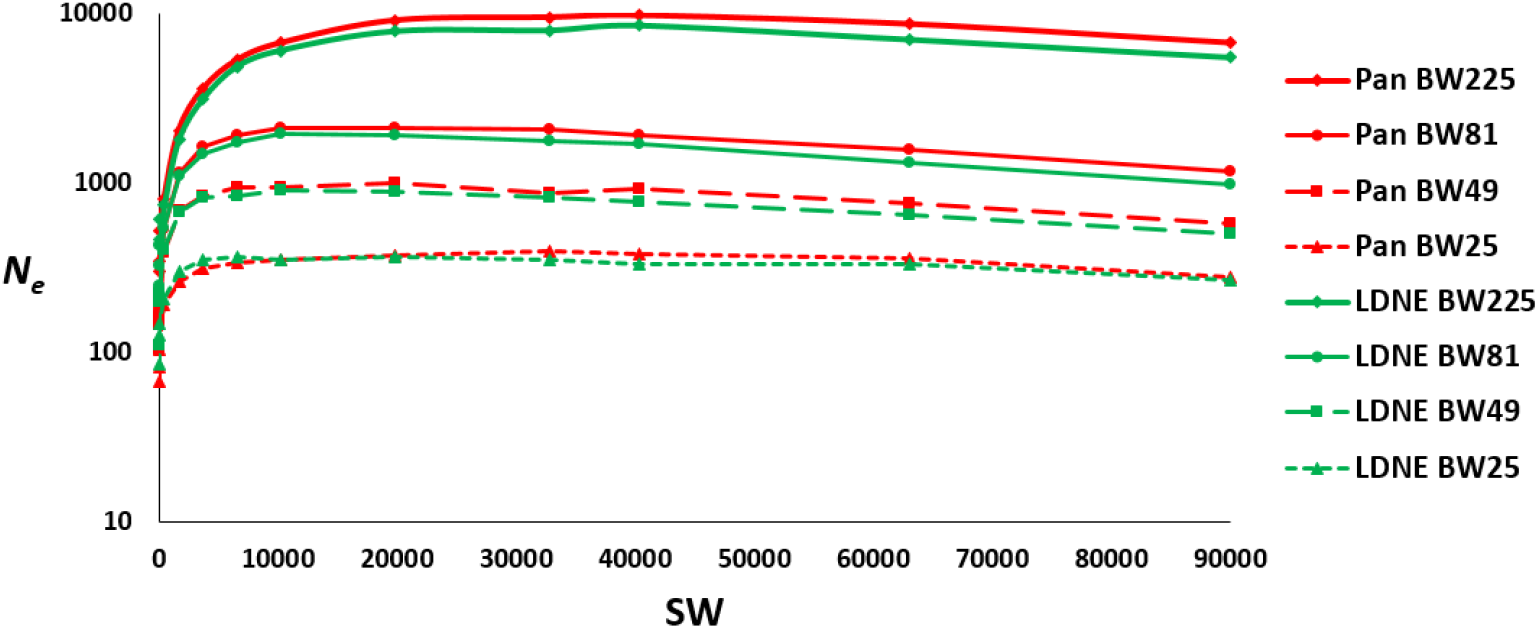
Estimates of *N*_*e*_ obtained with the software *currentNe2* (Santiago et al. 2025) assuming panmixia (red lines) and the software *LDNE* (Waples and Do 2008) (green lines), which also assumes panmixia. The different scenarios refer to different breeding windows (BW).

